# Long-term combination therapy with Metformin and Oxymetholone in a Fanconi Anemia mouse model

**DOI:** 10.1101/2023.08.16.553572

**Authors:** Craig Dorrell, Alexander M Peters, Qingshuo Zhang, Niveditha Balaji, Kevin Baradar, Makiko Mochizuki-Kashio, Angela Major, Milton Finegold, Chih-Wei Liu, Kun Lu, Markus Grompe

## Abstract

Fanconi Anemia (FA) is a disease caused by defective DNA repair which manifests as bone marrow failure, cancer predisposition, and developmental defects. Mice containing inactivating mutations in one or more genes in the FA pathway partially mimic the human disease. We previously reported that monotherapy with either metformin (MET) or oxymetholone (OXM) improved peripheral blood (PB) counts and the number and functionality of bone marrow (BM) hematopoietic stem progenitor cells (HSPCs) number in *Fancd2*^-/-^ mice. To evaluate whether the combination treatment of these drugs has a synergistic effect to prevent bone marrow failure in FA, we treated cohorts of *Fancd2*^-/-^ mice and wild-type controls with either MET alone, OXM alone, MET+OXM or placebo diet. Both male and female mice were treated from age 3 weeks to 18 months. The OXM treated animals showed modest improvements in blood parameters including platelet count (p=0.01) and hemoglobin levels (p<0.05). In addition, the percentage of quiescent HSC (LSK) was significantly increased (p=0.001) by long-term treatment with MET alone. However, the absolute number of progenitors, measured by LSK frequency or CFU-S, was not significantly altered by MET therapy. The combination of metformin and oxymetholone did not result in a significant synergistic effect on any parameter. Male animals on MET+OXM or MET alone were significantly leaner than controls at 18 months, regardless of genotype. Gene expression analysis of liver tissue from these animals showed that some of the expression changes caused by *Fancd2* deletion were partially normalized by metformin treatment. Importantly, no adverse effects of the individual or combination therapies were observed, despite the long-term administration.

**Highlights:**

- Long-term coadministration of metformin in combination oxymetholone is well tolerated by *Fancd2*^-/-^ mice.
- HSC quiescence in mutant mice was enhanced by treatment with metformin alone.
- Metformin treatment caused a partial normalization of gene expression in the livers of mutant mice.

## Introduction

Fanconi Anemia (FA) is a genetic disorder characterized by progressive bone marrow failure and an increased risk of cancer ^1,2^. The disease is caused by a mutation in any of the 21+ Fanconi DNA repair genes, which encode proteins responsible for the elimination of inter-strand DNA crosslinks. The disruption of this repair pathway causes the accumulation of DNA damage, which impairs replication and leads to the accelerated turnover of hematopoietic progenitor/stem cells and facilitates cancer-causing mutations.

Mouse models of this disease, created by knocking out the murine equivalent of each “Fanconi gene”, can replicate some of the disease phenotypes observed in human patients. *Fancd2*^-/-^ mice on the 129/S4 strain background are sterile due to hypogonadism, exhibit sensitivity to DNA crosslinking agents, and have an elevated frequency of epithelial tumors ^3^. They do not generally die of bone marrow failure but exhibit progressive hematopoietic disruptions including thrombocytopenia and a significant loss of progenitor/stem cells ^4^. This animal model has been used to evaluate the effects of a number of potentially therapeutic agents ^4,5^, including oxymetholone ^6^ and metformin ^7^. Oxymetholone is an androgen clinically approved for the treatment of anemia in FA patients by stimulating hematopoietic proliferation ^8,9^. In mice, long-term oxymetholone treatment was shown to significantly increase the frequency of WBC in the periphery and to stimulate the proliferation of BM progenitors ^6^. Metformin is a biguanide widely used to reduce blood sugar in type 2 Diabetes patients by suppressing hepatic glucose production via AMPK activation ^10^. Because both AMPK activation ^11^ and the radical scavenging abilities of biguanides are thought to be beneficial for the preservation of hematopoietic function, metformin was also tested in *Fancd2*^-/-^ 129/S4 mice. This study found that six months of metformin treatment reduced thrombocytopenia and significantly increased the frequency of hematopoietic progenitors ^7^. Metformin and oxymetholone clearly have different mechanisms of action in FA mice. Oxymetholone accelerated the cell cycle ^6^ of stem cells, likely aiding in their expansion. Conversely, metformin resulted in increased quiescence of stem cells ^7^, a finding that may reflect partial protection from DNA damage. A short-term clinical trial with metformin monotherapy has been performed in human FA patients and displayed hints of efficacy ^12^. However, androgen therapy which currently is the standard of care in FA was an exclusion criterion in our metformin study ^12^. To date a combination of both drugs has never been tested in either preclinical models or human trials. Given the clearly different mechanisms of action, we wished to determine here whether combination therapy was synergistically beneficial and safe.

Here we describe the results of long-term combined administration of oxymetholone and metformin to *Fancd2*^-/-^ 129/S4 mice determine whether this would yield stronger hematopoietic benefits. We observed modest improvements in platelet number (p=0.01) and hemoglobin levels (p<0.05) in OXM-treated animals and the frequency of quiescent HSC (LSK) was significantly increased (p=0.001) by long-term MET treatment. The total number of progenitors, measured by LSK frequency or CFU-S, was not significantly affected by drug treatment. Liver gene expression was partially normalized by MET treatment, including the reduced expression of kinetochore and other genes associated with mitosis. Although the combination of these two agents did not yield a dramatic improvement in murine hematopoiesis, they were well tolerated and may synergize more effectively in human patients.

## METHODS

### Mice

*Fancd2*^*-/-*^ mice (with a 129/S4 strain background) were bred as heterozygotes in our animal facility at Oregon Health & Science University. The metformin (3.75 g/kg), oxymetholone (0.3 g/kg), and metformin + oxymetholone (3.75 g/kg + 0.3 g/kg respectively) diets were custom milled with standard rodent diet (Purina Chow 5001) by Bio-Serv (Flemington, NJ) and administered to mice after weaning (3-4 weeks of age). All animals were treated in accordance with the guidelines of the Institutional Animal Care and Use Committee, protocol T_000000446.

### Blood analyses

Peripheral blood (50 µl) was collected by retroorbital sampling in EDTA-coated capillary tubes and complete blood counts (CBCs) were measured on a Hemavet 950FS Multi-species Hematology System (Drew Scientific, Inc, Dallas, TX).

### CFU-S progenitor assay

The colony-forming unit–spleen (CFU-S) assay was performed as described previously ^7^. Briefly, femoral BM was collected from mice at the time of sacrifice (18 months). These cells were retro-orbitally transplanted (1×10^6^ MNC each) into 3-4 wildtype 129/S4 recipient animals 24h after whole body x-ray irradiation. Twelve days after transplantation, the recipients were sacrificed and the number of macroscopic colonies on their spleens was scored.

### Flow cytometry

Femoral bone marrow cells were isolated (as described above) and labeled with a “Lin+” antibody cocktail (anti-CD3e, -CD4, -CD5, -CD8a, -B220, -Ter119, -NK1.1, -Mac1, and -Gr1), anti-c-kit, anti-Sca1, ki-67 (eBioscience, San Diego, CA) and Hoechst 33342 (Invitrogen). Acquisitions were performed on a FACSCanto II (BD Biosciences, San Jose, CA) equipped with blue, red, and UV lasers and analyzed using Flowjo (FlowJo, LLC).

### RNA-seq

RNA extraction, library construction, and sequencing of liver tissue samples were performed by Genewiz (South Plainfield, NJ). Read mapping and quantification was performed by the OPRC sequencing core. The resulting gene lists were analyzed using Ingenuity Pathway Analysis (Qiagen Silicon Valley, Redwood City CA), REACTome (reactome.org)^13^ and relationships were visualized with STRING (string-db.org) ^14^

### DNA adduct analysis

The experimental procedures for the extraction and enzymatic hydrolysis of tissue genomic DNA were based on previously described methods. ^15-17^ Liver tissues (∼20 mg) were homogenized in G2 solution from NucleoBond buffer kit by a stainless steel bead via the mechanical disruption of TissueLyzer (QIAGEN) for 10min (50 Hz). NucleoBond AXG 20 columns and NucleoBond buffer kit were purchased from Macherey-Nagel (Bethlehem, PA). Nanosep Centrifugal Devices (MWCO 3K) and stainless steel beads (5 mm) were purchased from Pall Lift Sciences (Port Washington, NY) and QIAGEN (Germantown, MD), respectively. Those biological samples in G2 solution were used for DNA purification according to the manufacturer instruction for NucleoBond AXG 20 sample preparation. The purified DNA was quantified by Nanodrop 2000 (Thermo Fisher Scientific). The extracted DNA (10 μg) was reduced by incubation with NaBH3CN (50 mM) and sodium phosphate buffer (100 mM, pH 7.2) for 17 h at 37 °C with gentle shake before enzymatic digestion. The DNA solution was added with 200 μL of 50 mM sodium phosphate/20 mM MgCl_2_ buffer (pH 7.2) along with known amounts of the internal standards ([13C1015N5]-N2-methyl-dG and [15N5]-N2-ethyl-dG) before digestion by DNase I (200 units), alkaline phosphatase (5 units), and phosphodiesterase (0.005 units) for 1 h at 37 °C. Following digestion, hydrolyzed DNA was filtered with the NanoSep 3 kDa filter at 8,000 rpm for 50 min to remove enzymes prior to HPLC purification and targeted nano-LC-ESI-MS/MS analysis for absolute quantification of DNA adducts^15-17^.

### Serum metformin measurement

A cohort of five 129/S4 animals were given metformin-containing food for a week. Serum was then isolated from retroorbitally collected blood and samples were submitted to NMS Labs (Horsham, PA) for analysis by high performance liquid chromatography / tandem mass spectrometry (LC-MS).

### Statistics

Prism (Graphpad, San Diego, CA) was used to perform 2-tailed, unpaired Student’s t tests for significance determination. Values of p < .05 were considered significant.

## RESULTS

### Long term administration of metformin and oxymetholone is well tolerated by Fancd2^-/-^ mice

To determine whether the combined, long-term administration of metformin and oxymetholone would provide a greater normalization of hematopoiesis than had been shown for each drug individually, a cohort of 162 mice was developed. These were used to populate sixteen groups, distinguished by genotype (*Fancd2*^-/-^ / WT), sex (M / F), and treatment (control / metformin / oxymetholone / metformin + oxymetholone). Of these, 89% (144/162) survived for the planned 18 month study duration and contributed tissues for the analyses. During monitoring, we observed that male mice in some of the drug treatment groups appeared leaner, and terminal weight measurements bore this out (**Supplementary Fig. 1a**). The effect of oxymetholone treatment alone was not significant, but metformin or dual treatment produced a significant weight reduction in both *Fancd2*^-/-^ and WT animals of approximately 23%. Neither drug significantly affected the weight of female mice at 18 months (**Supplementary Fig. 1b**). Liver tissue was collected for histological study and scored for steatosis (**Supplementary Fig. 2a**) and DNA adduct frequency (**Supplementary Fig. 2b**), but this revealed only a modest (non-significant) increase in DNA damage in the liver cells of mutant mice.

### Peripheral blood monitoring revealed modest normalization of peripheral blood characteristics

A previous study showed that at 18 months, *Fancd2*^-/-^ mice exhibited peripheral pancytopenia as well as a significant disruptions of hemoglobin levels and MCV ^4^. In the new cohort used here, however, the mutant animals had only mild peripheral blood deficits but almost none that were statistically significant (**Fig.1 a-g**). By performing sex-specific analysis we able to identify a significant deficit; at 18 months. Female *Fancd2*^-/-^ mice had a 23% reduction in platelets (**Fig. 1g**; p=0.02). In female mutants treated with oxymetholone this mild platelet deficit was eliminated (p=0.01), and mutant mice of both sexes had significantly increased hemoglobin levels after oxymetholone or combination treatment (**Fig.1d**; p<0.05). The blood levels of metformin were analyzed and confirmed to be in the therapeutic range of about 15 µM (Supplementary Fig.3).

**Figure 1.**
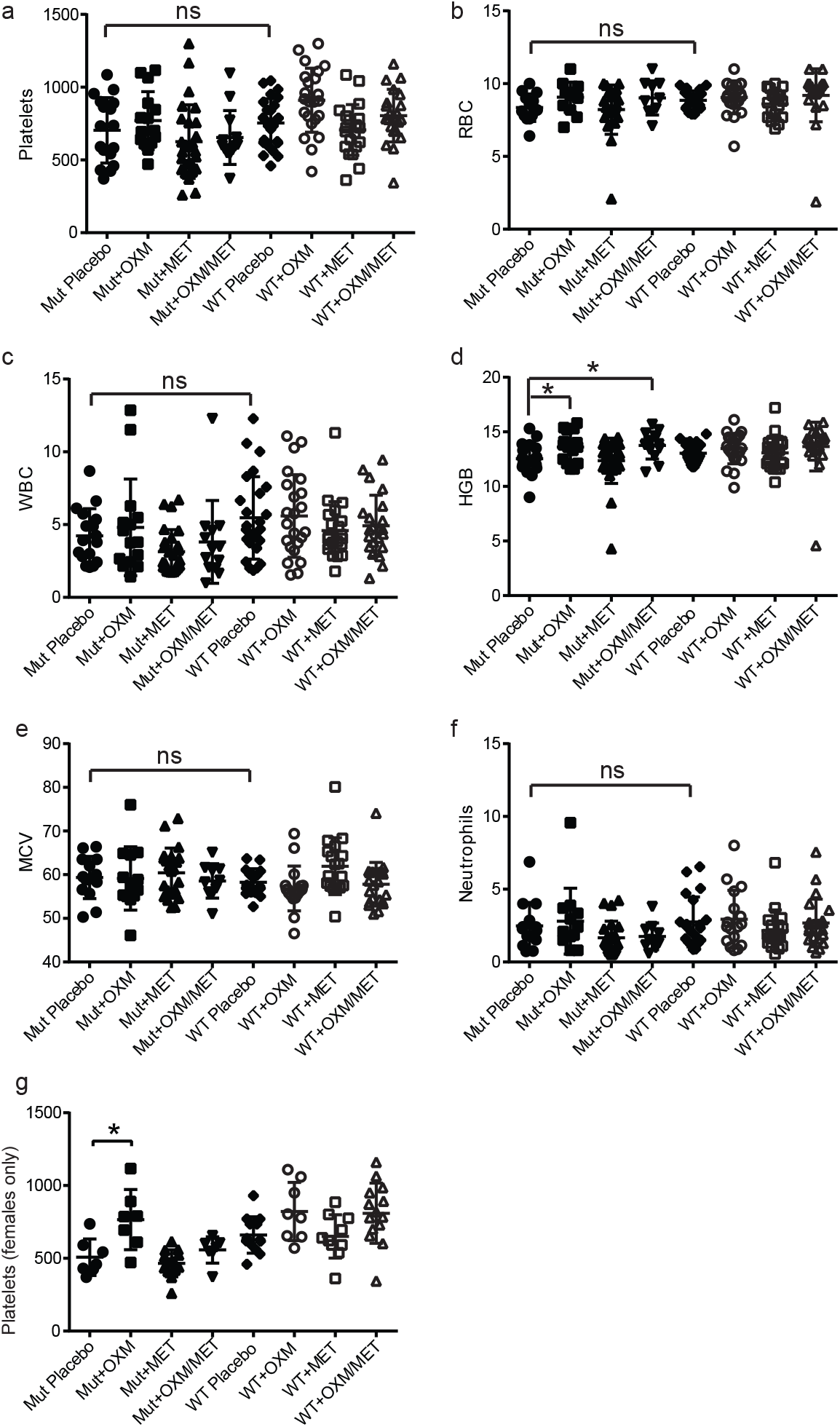
Most peripheral blood parameters were not significantly affected by long term drug treatment. After 18 months, no significant alterations of platelet count (a), red blood cell frequency (b), white blood cell frequency (c), mean corpuscular volume (e), or neutrophil frequency (f) were observed. Oxymetholone treatment or metformin + oxymetholone treatment yielded a modest increase in hemoglobin levels (d) and platelet frequency was normalized (p=0.015) in female mutants (g).

### Hematopoietic progenitor quiescence, but not frequency, was enhanced by metformin

Femoral bone marrow was collected at 18 months and used to survey the progenitor compartment. The *Fancd2*^-/-^ animals had a LSK cell deficit (34% of wild-type ;p=0.001) deficit which was not significantly ameliorated by any of the treatments (**Fig. 2a**). Mutants also had a significantly higher number of proliferating LSK, measured either by >2n DNA content (23% [p=0.001]; **Fig. 2b**) or G0 frequency (19% [p<1×10^−4^]; **Fig. 2c**). Metformin treatment reversed this deficit. For a functional assessment of stem/progenitor frequency in BM, d12 CFU-S assays were also performed (**Fig. 2d**). *Fancd2*-/-mice had a 62% (p=0.001) lower CFU-S colony frequency than wild-type controls, which was not significantly affected by drug treatment.

**Figure 2.**
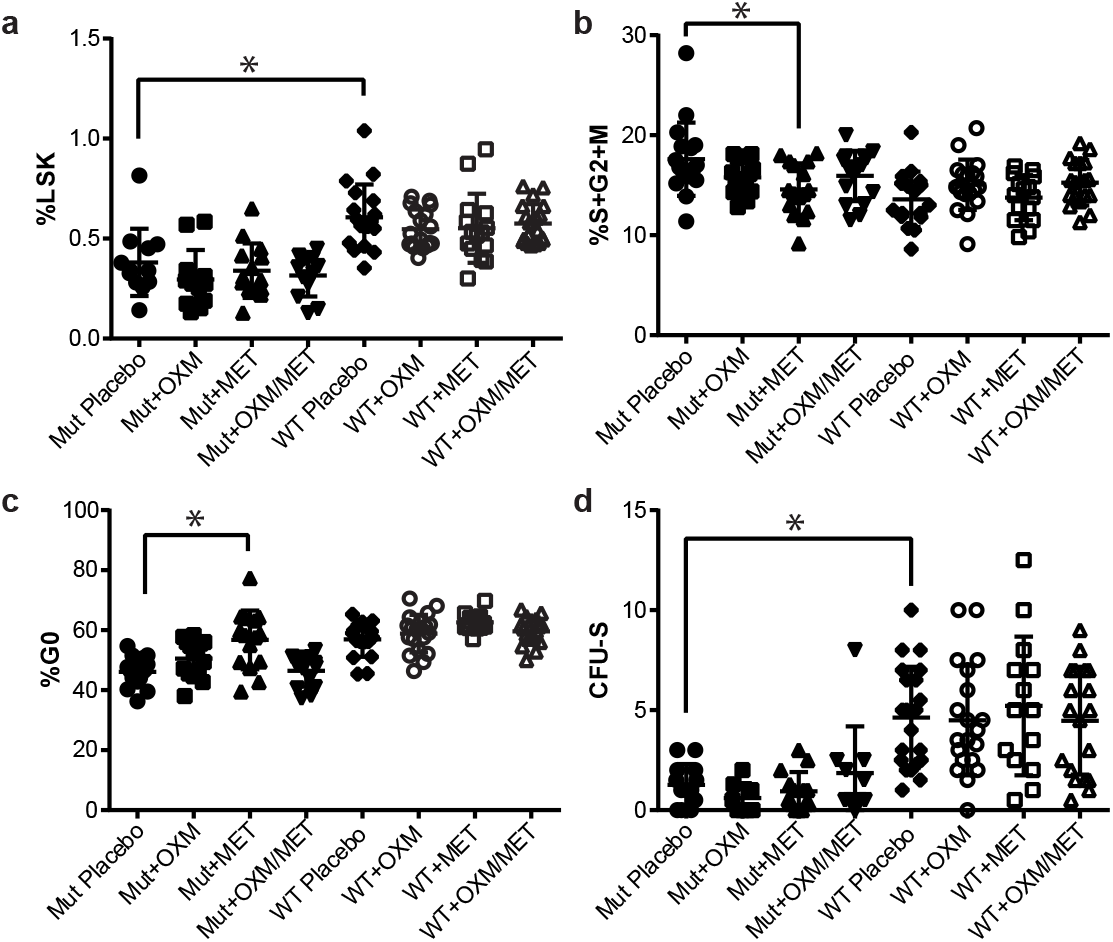
Progenitor quiescence but not frequency was increased by long-term metformin treatment. The frequency of Lin^-^Sca^+^c-Kit^+^ (KSL) progenitors in femoral bone marrow of **Fancd2**^-/-^ mice was reduced relative to wildtype controls (p=0.001) but drug treatment did not significantly change this frequency in either genotype (a). Measurements of KSL quiescence including DNA content and ki-67 analysis showed that metformin treatment significantly decreased the frequency of proliferating progenitors (b) and increased the percentage in G0 (c). The frequency of d12 CFU-S was found to be significantly (p<0.001) reduced in mutant animals but this was not corrected by drug treatment.

**Figure 3.**
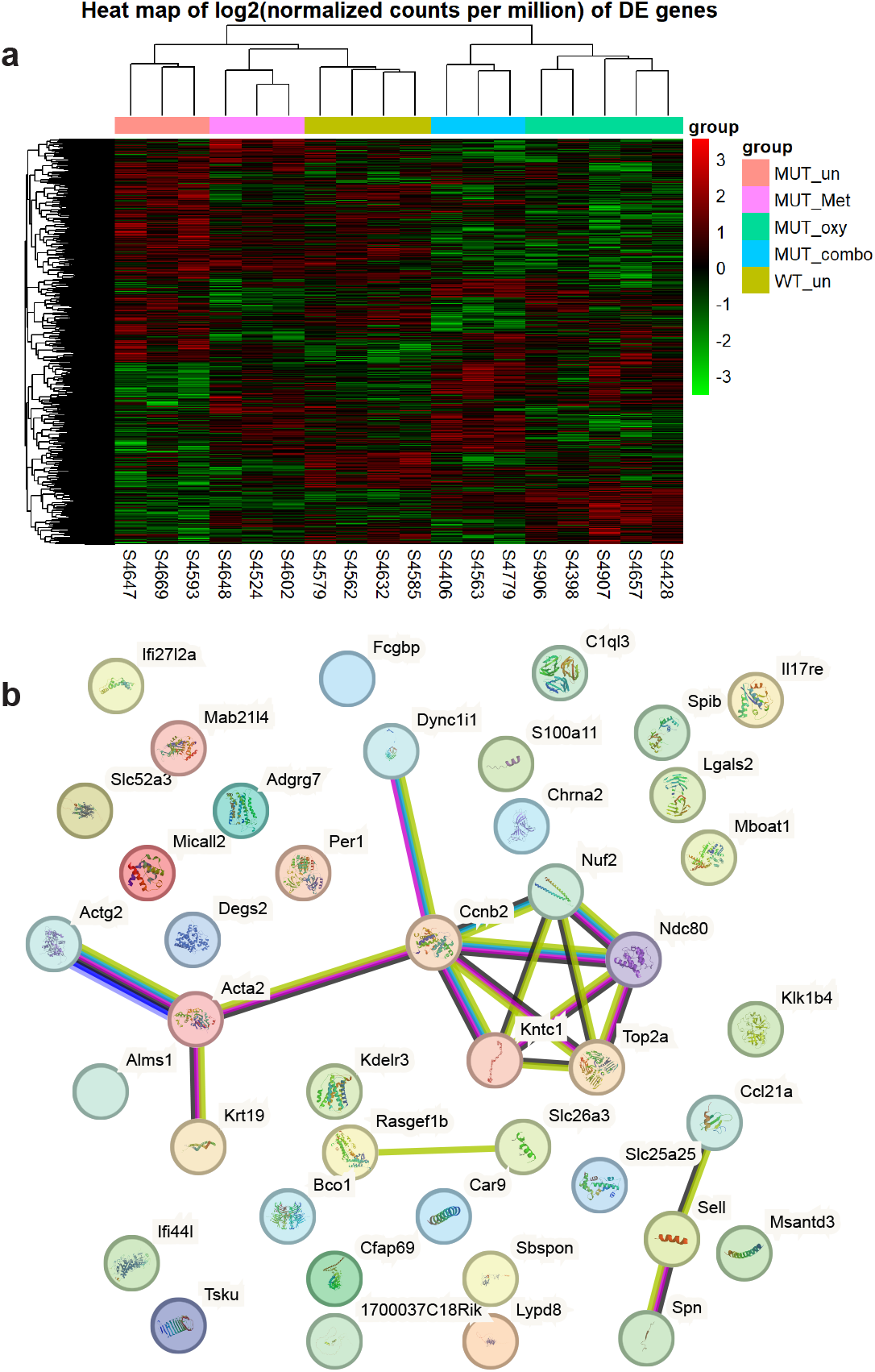
Gene expression in liver is altered by drug treatment. (a) Comparative gene expression with unsupervised gene clustering after removing surrogate variable effects. Note that the metformin treated mutants cluster more closely to wild-type than the untreated mutants. Similarly, combined treatment of metformin and oxymetholone clustered closer to wild-type than oxymetholone alone. (b) STRING analysis of mutant liver-expressed genes significantly normalized by metformin treatment. Most of the normalized gene expression events were unconnected, but the co-normalization of five different kinetochore component-encoding genes was detected.

### Metformin partially normalized liver gene expression in Fancd2^-/-^ mice

To seek subtle alterations in gene expression and assess whether drug treatment might reverse them, RNA was collected from twenty livers sampled from control-treated male WT and all four treatment subgroups of male mutant mice. RNA-seq analysis showed that a total of 278 genes were significantly (p<0.05) differentially (≥2x change) expressed between untreated WT and *Fancd2*^-/-^ animals, indicating the effects of Fancd2 loss on liver gene expression. Of these, 51 were found to be also significantly differentially expressed between control- and metformin-treated *Fancd2*^-/-^ liver (**Table 1**). Interestingly, in each case the effect of metformin was corrective, i.e. the metformin-treated group’s gene expression moved away from its state in control *Fancd2*^-/-^ mice towards its state in control WT mice. With String analysis (**Supplemental Fig. 3**), which illustrates reported relationships between individual genes, the most striking observation is that five of the genes (Ccnb2, Ndc80, Nuf2, Top2a, and Kntc1) that were downregulated in control *Fancd2*^-/-^ but not metformin-treated *Fancd2*^-/-^ mice are involved in kinetochore formation and cell cycling. REACTome analysis (**Table 2**) yields a similar observation; nearly all of the significantly associated pathways were cell cycle related.

**Table 1.** Genes differentially expressed between control- and metformin-treated Fancd2-/-liver. Expression levels determined by RNA-seq were analyzed using “limma voom” workflow; the listed genes were significantly (p<0.05) differentially expressed.

**Table 2.** Pathways differentially expressed between control- and metformin-treated Fancd2-/-liver. REACTOME analysis of the differentially expressed genes shows an association with multiple pathways related to cell division.

## DISCUSSION

The use of androgens such as oxymetholone to improve erythropoiesis and thrombopoiesis in Fanconi anemia patients is well established ^8,9^, and in some cases is sufficient to prevent bone marrow failure. Even in cases where hematopoietic stem cell transplantation (HSCT) is strongly indicated, treatment with androgens can act as a bridging therapy until a transplant can be arranged ^2,18^. Consequently, it is useful to know whether newer therapeutic options such as metformin can be safely and effectively combined with androgens in the treatment of FA.

Our study was designed with the assumption that the hematopoietic phenotypes of Fancd2 mice would be similar to those observed in earlier studies from our group using this strain ^4,6^. We previously performed terminal analyses at around 6 months of age ^7^ and therefore hypothesized that an 18 month study would result in progression of the hematopoietic phenotype and yield an even more severe anemia. This, however, did not prove to be correct. Although the experimental animals examined here were derived from the same breeding colony as previously used, their hematopoietic phenotype was very mild, especially in the peripheral blood. The precise cause of this change is not known, but others have demonstrated that environmental and microbiome influences can cause phenotypic drifts ^19,20^. The lack of a strong phenotype in peripheral blood made the assessment of therapeutic benefit difficult. Whereas a six month course of oxymetholone was previously reported to significantly improve platelet, RBC, hemoglobin, MCV, and hematocrit in *Fancd2*^-/-^ mice^6^, we observed only a small improvement in hemoglobin and a correction of thrombocytopenia that was exclusive to the female subgroup. A similar study employing metformin treatment ^7^ described enhancements of WBC, hemoglobin, and platelet counts which we did not observe despite a much longer treatment duration. Nonetheless, the previously reported deficits in the stem cell compartment were replicated here. *Fancd2*^-/-^ mice have significantly fewer bone marrow stem cells as measured by immunophenotyping and the CFU-S assay. In addition, the percentage of quiescent LSK stem cells was clearly decreased in mutant animals. Interestingly, the changes observed at age 18 months here were not more severe than previously seen at the 6 month time-point. This suggests that the bone marrow failure in *Fancd2*^-/-^ mice is not progressive with time. None of the treatment regimens significantly improved the number of stem cells, but metformin – as reported earlier – significantly enhanced stem cells quiescence. Overall, it appears that oxymetholone and metformin did not synergize to rescue the stem cell number in this model. Importantly, however, no adverse effects were observed either with the combination treatment.

Given the important metabolic effects of androgens and the AMPK activator metformin, we wished to assess the state of the liver in mutants and drug treated animals. Although liver histology showed no consistent abnormalities, including in the degree of steatosis, the gene expression profile in the livers of *Fancd2*^-/-^ mice showed many dysregulations. Interestingly, treatment with metformin caused significant normalization of the expression of 52 FA dysregulated genes in the liver. Pathways including kinetochore metaphase signaling (p=4×10^−5^) and DNA double-strand break repair by homologous recombination (p=8.1×10^−4^) were beneficially impacted by metformin, but not oxymetholone. Metformin is thought to have multiple chemical interactions and the mechanism by which it might support mitosis or DSBR is not clear, but if it reduces overall frequency of DNA damage – perhaps by aldehyde scavenging – both of those processes might be more efficient.

Perhaps the most important conclusion of our study is that an 18 month co-administration of oxymetholone and metformin was well tolerated by *Fancd2*^-/-^ mice. None of the hematopoietic endpoints assessed was adversely affected. In addition, survival to the experimental endpoint was identical to that of the other treatment groups and no additional pathology was observed at necropsy.

Metformin monotherapy in human FA patients has recently demonstrated potential benefit ^12^. Our preclinical data here provide support for the inclusion of androgen treated FA patients in future clinical trials of metformin in Fanconi Anemia.

## Acknowledgements

We greatly appreciate the assistance of the veterinary and technical staff of the OHSU Department of Comparative Medicine for the monitoring and husbandry of the large cohort of mice used in this study. In addition, we are grateful to the staff of the OHSU flow cytometry core for their assistance and the use of key analytical equipment. RNA quality assessment, library assembly, and RNA-seq was provided by Suzanne Fei, Lina Gao, and Karina Ray of the Oregon National Primate Research Center.

These studies were supported by a 2016-2021 program project grant from the NIH 5P01HL048546-24.

## Figure Legends

**Supplementary Figure 1** - Metformin treatment reduced the weight of male mice. Mice were weighed at 18 months as part of the terminal harvest. Regardless of genotype, male animals (a) fed metformin (p=0.01) or metformin + oxymetholone (p=0.03) supplemented diet were significantly leaner than the control group. No significant effect was observed in female animals (b).

**Supplementary Figure 2** – Analysis of liver quality at 18 months. Histological analysis of liver sections for the incidence and grade of steatosis showed no significant correlations between genotype or drug treatment (a). The frequencies of ethyl and methyl dG DNA adducts trended higher in mutant liver tissue than in controls, but this was not significant and was not significantly altered by metformin treatment (b).

**Supplementary Figure 3** – Serum metformin concentrations in mice fed metformin-containing diet. LC-MS measurements of serum from five animals after seven days of diet consumption are shown.

