## Supplementary figures and images for "Long-term combination therapy with Metformin and Oxymetholone in a Fanconi Anemia mouse model"

### Supplemental Figure 1

#### Supplementary Fig. 1

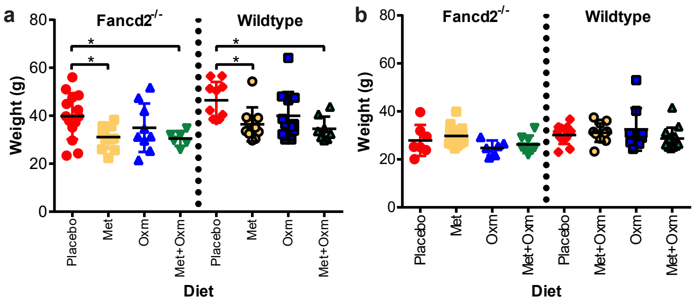

### Supplemental Figure 2

Supplementary Fig. 2

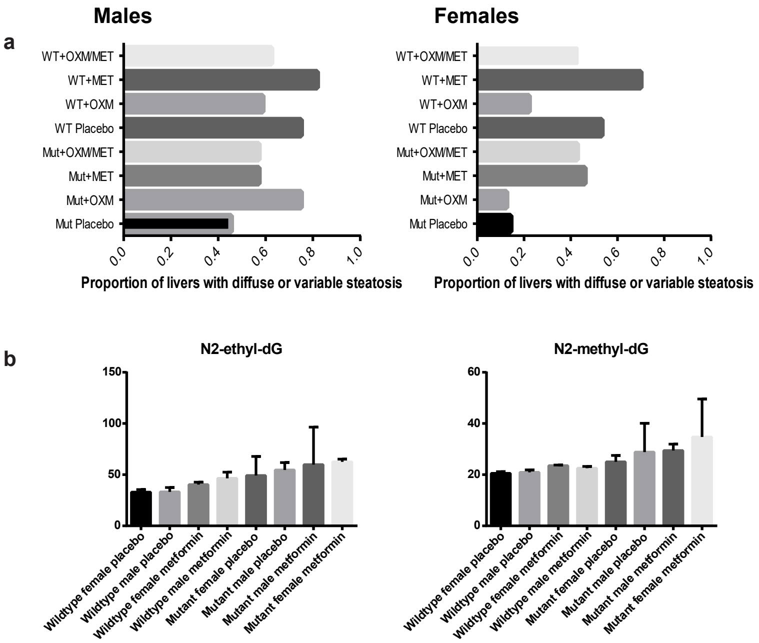

### Supplemental Figure 3

Supplementary Fig. 3

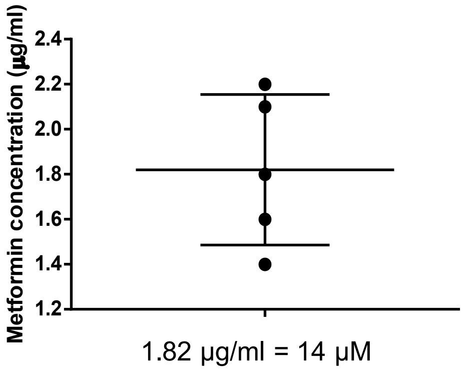


1.82 µg/ml = 14 µM

**d**
