## Supplemental Table 1 for "Long-term combination therapy with Metformin and Oxymetholone in a Fanconi Anemia mouse model"

| Gene | D2-/- MET vs D2-/- ctrl (Log2FC) | p | FDRp | D2-/- ctrl vs WT ctrl (Log2FC) | p | FDRp |
|---|---|---|---|---|---|---|
| Adgrg7 | -3.08 | 0.01 | 1.00 | 5.01 | 0.00 | 0.16 |
| Slc26a3 | -3.59 | 0.00 | 1.00 | 3.91 | 0.00 | 0.46 |
| Lypd8 | -2.34 | 0.03 | 1.00 | 3.34 | 0.00 | 0.66 |
| Krt19 | -3.17 | 0.00 | 0.25 | 2.77 | 0.00 | 0.08 |
| Car9 | -2.34 | 0.04 | 1.00 | 2.43 | 0.01 | 0.87 |
| Actg2 | -3.80 | 0.01 | 1.00 | 2.35 | 0.02 | 0.87 |
| Lgals2 | -2.60 | 0.05 | 1.00 | 2.16 | 0.03 | 0.88 |
| Acta2 | -2.30 | 0.02 | 1.00 | 2.06 | 0.01 | 0.87 |
| Slc52a3 | -2.00 | 0.05 | 1.00 | 2.03 | 0.02 | 0.87 |
| Ccl21a | -3.26 | 0.01 | 1.00 | 1.96 | 0.02 | 0.87 |
| Mboat1 | -1.91 | 0.02 | 1.00 | 1.90 | 0.00 | 0.86 |
| Degs2 | -1.97 | 0.03 | 1.00 | 1.83 | 0.01 | 0.87 |
| Gm15348 | -2.87 | 0.01 | 1.00 | 1.81 | 0.01 | 0.87 |
| Kdelr3 | -2.17 | 0.00 | 1.00 | 1.66 | 0.00 | 0.86 |
| Gm37736 | -3.11 | 0.01 | 1.00 | 1.64 | 0.04 | 0.88 |
| Cfap69 | -1.76 | 0.03 | 1.00 | 1.63 | 0.01 | 0.87 |
| Il17re | -1.73 | 0.05 | 1.00 | 1.47 | 0.04 | 0.88 |
| Gm9320 | -1.29 | 0.05 | 1.00 | 1.47 | 0.00 | 0.87 |
| C1ql3 | -1.68 | 0.04 | 1.00 | 1.42 | 0.02 | 0.87 |
| Mab21l4 | -1.96 | 0.02 | 1.00 | 1.31 | 0.04 | 0.88 |
| 1700037C18Rik | -1.16 | 0.03 | 1.00 | 1.28 | 0.00 | 0.86 |
| Bco1 | -1.31 | 0.02 | 1.00 | 1.23 | 0.00 | 0.87 |
| S100a11 | -0.85 | 0.05 | 1.00 | 1.18 | 0.00 | 0.76 |
| Klk1b4 | -1.18 | 0.01 | 1.00 | 1.08 | 0.00 | 0.76 |
| Fcgbp | -1.43 | 0.02 | 1.00 | 1.01 | 0.03 | 0.88 |
| Chrna2 | 1.88 | 0.00 | 1.00 | -1.01 | 0.04 | 0.88 |
| Alms1 | 1.53 | 0.01 | 1.00 | -1.04 | 0.03 | 0.88 |
| Top2a | 1.93 | 0.00 | 1.00 | -1.16 | 0.01 | 0.87 |
| Dync1i1 | 1.60 | 0.02 | 1.00 | -1.20 | 0.03 | 0.88 |
| Micall2 | 1.16 | 0.03 | 1.00 | -1.28 | 0.01 | 0.87 |
| Ifi44l | 1.36 | 0.02 | 1.00 | -1.28 | 0.01 | 0.87 |
| Rasgef1b | 0.93 | 0.03 | 1.00 | -1.28 | 0.00 | 0.75 |
| Ifi27l2a | 1.41 | 0.04 | 1.00 | -1.39 | 0.02 | 0.87 |
| Ccnb2 | 1.51 | 0.02 | 1.00 | -1.43 | 0.02 | 0.87 |
| Sell | 1.13 | 0.03 | 1.00 | -1.46 | 0.00 | 0.76 |
| Per1 | 2.20 | 0.00 | 1.00 | -1.47 | 0.01 | 0.87 |
| Msantd3 | 2.79 | 0.00 | 1.00 | -1.49 | 0.03 | 0.87 |
| Slc25a25 | 1.62 | 0.03 | 1.00 | -1.53 | 0.01 | 0.87 |
| Nuf2 | 2.20 | 0.01 | 1.00 | -1.56 | 0.05 | 0.88 |
| Gm29737 | 2.47 | 0.00 | 1.00 | -1.61 | 0.01 | 0.87 |
| Gm18119 | 1.35 | 0.04 | 1.00 | -1.62 | 0.01 | 0.87 |
| Gm6831 | 2.02 | 0.04 | 1.00 | -1.66 | 0.04 | 0.88 |
| Gm12473 | 1.62 | 0.03 | 1.00 | -1.68 | 0.02 | 0.87 |
| Spn | 1.37 | 0.01 | 1.00 | -1.90 | 0.00 | 0.46 |
| Sbspon | 2.38 | 0.03 | 1.00 | -2.02 | 0.04 | 0.88 |
| Spib | 1.98 | 0.02 | 1.00 | -2.04 | 0.01 | 0.87 |
| Kntc1 | 1.82 | 0.03 | 1.00 | -2.05 | 0.01 | 0.87 |

| | | | | | | |
|---|---|---|---|---|---|---|
| Tsku | 2.46 | 0.00 | 1.00 | -2.05 | 0.00 | 0.86 |
| Gm50022 | 1.68 | 0.01 | 1.00 | -2.27 | 0.00 | 0.62 |
| Ndc80 | 3.34 | 0.00 | 1.00 | -2.50 | 0.00 | 0.86 |
| Gm4468 | 2.20 | 0.01 | 1.00 | -2.62 | 0.00 | 0.75 |
