## Supplemental Table 2 for "Long-term combination therapy with Metformin and Oxymetholone in a Fanconi Anemia mouse model"

| Pathway name | Entities found | Total | Ratio | p value | FDR |
|---|---|---|---|---|---|
| Mitotic Prometaphase | 6 | 211 | 0.01 | 0.00 | 0.01 |
| Resolution of Sister Chromatid Cohesion | 5 | 134 | 0.01 | 0.00 | 0.01 |
| Cell Cycle, Mitotic | 9 | 596 | 0.04 | 0.00 | 0.02 |
| Amplification of signal from unattached kinetochores via a MAD2 inhibitory signal | 4 | 94 | 0.01 | 0.00 | 0.02 |
| Amplification of signal from the kinetochores | 4 | 94 | 0.01 | 0.00 | 0.02 |
| Mitotic Spindle Checkpoint | 4 | 111 | 0.01 | 0.00 | 0.03 |
| EML4 and NUDC in mitotic spindle formation | 4 | 121 | 0.01 | 0.00 | 0.03 |
| Cell Cycle | 9 | 734 | 0.05 | 0.00 | 0.04 |
| RHO GTPases Activate Formins | 4 | 149 | 0.01 | 0.00 | 0.04 |
| Mitotic Anaphase | 5 | 249 | 0.02 | 0.00 | 0.04 |
| Mitotic Metaphase and Anaphase | 5 | 250 | 0.02 | 0.00 | 0.04 |
| Polo-like kinase mediated events | 2 | 23 | 0.00 | 0.00 | 0.05 |
| TLR3 deficiency - HSE | 1 | 1 | 0.00 | 0.00 | 0.05 |
| Transcription of E2F targets under negative control by DREAM complex | 2 | 25 | 0.00 | 0.00 | 0.05 |
| Cell Cycle Checkpoints | 5 | 279 | 0.02 | 0.00 | 0.05 |
| NOTCH4 Intracellular Domain Regulates Transcription | 2 | 26 | 0.00 | 0.00 | 0.05 |
| M Phase | 6 | 416 | 0.03 | 0.01 | 0.05 |
| Separation of Sister Chromatids | 4 | 195 | 0.01 | 0.01 | 0.06 |
